# Basal Role for Nrf2-transgne on transcriptional (mRNA/miRNA) regulation in the mouse myocardium

**DOI:** 10.1101/490953

**Authors:** Arun Jyothidasan, Gobinath Shanmugam, John Zhang, Brain Dally, David Crossman, Namakkal S. Rajasekaran

## Abstract

**Background:** NFE2L2 (nuclear factor, erythroid 2 like 2; Nrf2), is a stress-responsive transcription factor that regulates cellular redox homeostasis. The action mechanism of Nrf2 occurs through sensing oxidative stress in the cellular environment in responses to various stresses including altered metabolism, toxic insult and xenobiotic stresses. However, under the basal condition, the role of excess Nrf2 on global miRNA–mRNA interactions in the myocardium is unknown. Here, we tested a hypothesis that excess Nrf2 (transgene) will promote the transcription of antioxidants, which then might escalate the basal defense mechanisms in the heart. Furthermore, we investigated whether changes in the miRNA profile might have a strong role on the transcriptome in TG hearts.

**Methods:** Non-transgenic (NTG), and Cardiac specific Nrf2 transgenic (Nrf2-TG) mice in the C57/BL6 background at the age of 6-8 months were used to examine transcriptional and post-transcriptional signatures (mRNA and miRNA expressions) by performing next generation sequencing (NGS) for RNA (RNAseq) and small RNA (sRNAseq) for miRNA expressions in the heart (n=3-6/group). Validation of the NGS data was performed by quantitative real time PCR (qRT-PCR) using specific primers targeting selected miRNAs or mRNAs. Finally, in-silico analyses were performed to determine the miRNA–mRNA interactome networks and the pathways that are potentially regulated by these networks.

**Results:** NGS analysis for mRNA indicated that there were 5727 differently expressed genes (DEGs) in the Nrf2-TG vs. NTG myocardium. Of which, 3552 were upregulated and 2175 were downregulated significantly. Small RNAseq analysis revealed that 112 miRNAs were significantly altered in Nrf2-TG versus NTG hearts. Among these miRNAs, 68 were upregulated and 44 were downregulated significantly. Validation of key targets for mRNA and miRNA by qPCR confirmed the NGS results. In-silico analysis (IPA) revealed that the majority of the miRNAs and mRNAs that are significantly altered in response to transgene expression are potentially involved in Nrf2 mediated oxidative stress response, ILK Signaling, hypoxia signaling in the cardiovascular system and glutathione-mediated detoxification etc.

**Conclusion:** These high-throughput sequencing data sets from cardiac specific Nrf2-TG mice revealed the transcriptome-wide effects of Nrf2 in the myocardium. Furthermore, our comprehensive analysis indicates that Nrf2 may directly or indirectly regulates these sub-sets of cardiac miRNA-mRNA interactome networks under basal physiological setting.

## Introduction

Transcription factor, nuclear factor erythroid 2 like 2 (NFE2L2/Nrf2) regulates antioxidant and cytoprotective genes in response to stress (1–3). It is believed that under non-stress (basal physiological) conditions, Nrf2 seems to have minimal or no role (2, 4–6). It has also been reported that ablation of Nrf2 in mice (Nrf2^−/−^) did not evoke any obvious pathological processes until the middle age under unstressed conditions (2). However, Nrf2^−/−^ mice developed susceptibility to various stresses (7–10) (i.e. oxidative, exercise, environmental, chemical and surgical stress), leading to end organ dysfunction including pathological remodeling of myocardium. Normally, Kelch like ECH associated protein 1 (Keap1) binds with Nrf2 and suppresses its nuclear import by rapid ubiquitination and subsequent degradation by the 26S proteasome (2, 6, 11, 12). Thus far, many conditions such as oxidative, electrophilic stress, etc. (13) trigger the dissociation of Nrf2 from Keap1 and permits its nuclear translocation and transactivate its target genes.

Nrf2 is ubiquitously expressed in mammalian organs/cells (1–3). Several reports have revealed that Nrf2-antioxidant genes were down-regulated during aging and other oxidative conditions in mouse myocardium, skeletal muscle, rat liver and skeletal muscle of humans (14–16). Nrf2 has been known for its beneficial effects for several decades against oxidative stress diseases including cancer (17), cardiovascular diseases (18), inflammation(19), pulmonary disease (20), and neurodegeneration (21, 22). Previously we reported that loss of Nrf2 significantly affects the global transcriptome and expression of miRNAs in the mouse myocardium suggesting its minimal role in biological functions (3). However, when Nrf2 is expressed excessively in the myocardium (under unstressed conditions), its role on transcriptional regulation and subsequent implication in biological functions are largely unknown. Therefore, in order to test whether excessive Nrf2 in the myocardium can evoke its transcriptional role on its known and unidentified targets, we generated a heart-specific transgenic mouse with mNrf2.

Here, we have performed next generation sequencing (NGS) using RNA and miRNA from non-transgenic (NTG) and mNrf2-TG mice in the myocardium to determine the transcriptome and small-RNA profiles. This deep sequencing results indicated that the TG hearts showed 5727 genes significantly (>1.5 fold) altered when compared to NTG mice. Furthermore, small RNA sequencing revealed that 112 miRNAs were significantly changed in the TG myocardium. Ingenuity pathway analysis (IPA) was performed to elucidate the uncovered pathways and their regulatory networks arising from the excess Nrf2 in the myocardium. These analyses revealed that the topmost pathways altered in TG mice were Nrf2 mediated oxidative stress response, ILK Signaling, hypoxia signaling in the cardiovascular system, glutathione-mediated detoxification, and cardiac hypertrophy signaling. Overall, these findings will facilitate a better understanding on the uncovered pathways (mRNAs and miRNAs) that are potentially regulated by Nrf2.

## Methods

### Reagents

RNeasy kit (74106), reverse transcription kit (205313), and QuantiTect SYBR Green PCR kit (cat. 204145) were purchased from Qiagen Inc., Valencia, CA. Primers for qPCR were designed using Primer Bank website (https://pga.mgh.harvard.edu/primerbank/) and purchased from Integrated DNA Technologies, Coralville. All other chemicals including reduced glutathione, RNA*later* meta-phosphoric acid were purchased from Sigma-Aldrich unless otherwise stated.

### Animals

Cardiac specific Nrf2-transgenic (mNrf2-TG) and its littermate control non-transgenic (NTG) mice with C57/Bl6J background at 6 - 8 months of age were used. Mice were housed under controlled temperature and humidity, a 12 h light/dark cycle, and fed with a standard rodent diet and water *ad libitum*. The Institutional Animal Care and Use Committee (IACUC) at the University of Alabama at Birmingham and the University of Utah, Salt Lake City, Utah approved all animal experiments, in accordance with the standards established by the US Animal Welfare Act.

### Myocardial glutathione levels

Myocardial levels of reduced GSH were assessed by a GSH detection kit from Cayman (Ann Arbor, MI, USA). In brief, heart tissues (25mg) from NTG and Nrf2-TG mice was homogenized with MES buffer and centrifuged at 5000 rpm for 5 min at 4°C. An aliquot of the supernatant was stored for protein determination and equal amount of 10% meta- phosphoric acid (MPA) was added to the remaining samples to precipitate the proteins. MPA extracts were treated with triethanolamine (TEAM) and samples were mixed with 150μl of reaction mixture cocktail containing MES buffer, NADPH, Glutathione Reductase enzyme and DTNB and immediately the enzymatic-recycling assay was performed as per the manufacturer’s instruction using a plate reader. GSH standards were prepared and processed similarly to generate a standard graph (2).

### Isolation of RNA and real-time qPCR analysis

NTG, and Nrf2-TG mice heart tissues (20-30mg) preserved in RNA (n=4-6) were homogenized and total RNA was extracted using RNeasy mini kit (Qiagen, 74106). RNA (1.25 μg) was used to synthesize the cDNA using with QuantiTect reverse transcription kit (Qiagen, 205313). Quantitative RT-PCR was performed using 25-50ng cDNA and 1 pmol primer (Table 1) in a 10 μl SYBR green reaction mix (Qiagen, 204056) and amplified in a Roche Light Cycler 480 (Roche, Basel, Switzerland). Relative expression was quantified using Ct values, and expression fold-change was calculated by normalization to the Ct of housekeeping genes *Gapdh* or *Arbp1* according to the 2^−ΔΔCt^ methods (2, 9).

### Next generation mRNA and miRNA sequencing

Myocardial RNA was extracted from 6-8 months old male Nrf2 Transgenic and Non transgenic mice using RNeasy Mini Kit (Qiagen, Cat. 74,106) as per the manufacturer’s instructions. After confirming sample purity, intact poly(A) transcripts were purified from total RNA using oligo(dT) magnetic beads. Sequencing libraries for mRNA were prepared with the TruSeq Stranded mRNA Library Preparation Kit (Illumina, RS-122-2101, RS-122-2102). Purified libraries were qualified using D1000 ScreenTape assay (Agilent, Cat. 5067– 5582/3) with 2200 TapeStation Instrument (Agilent Technologies). The cBot was used to apply 18pM of the sequencing library to a TruSeq v3 flowcell (Illumina) and the TruSeq SR Cluster Kit (Illumina, Cat. GD-401-3001) was used for clonal amplification. Finally, the flowcell was transferred to the HiSeq 2000 instrument and used in 50 cycle single read sequence run performed with TruSeq SBS Kit v3-HS reagents (Illumina, Cat. FC-401-3002). Novoindex (2.8) was used to create a reference index on a combination of hg19 chromosome and splice junction sequences. Splice junction sequences were generated with USeq (v8.6.4) MakeTranscriptome using Ensembl transcript annotations (build 67). Reads were aligned to the transcriptome reference index described above using Novoalign (v2.08.01), allowing up to 50 alignments for each read. USeq’s SamTranscriptomeParser application was used to select the best alignment for each read and convert the coordinates of reads aligning to splices back to genomic space. Differential gene expression was measured using USeq’s DefinedRegionDifferentialSeq application. The number of reads aligned to each gene was calculated. The counts were then used in DESeq (v1.24.0), which normalizes the signal and determines differential expression.

Similarly, mature miRNA transcripts were isolated from 6 to 8 months old Nrf2-TG (n = 3/group) myocardium using the miRNeasy Kit (Qiagen, Cat. 217,004). The sample purity was first confirmed. To prepare small RNA sequencing libraries, NEBNext Multiplex Small RNA Library Prep Set for Illumina (NEB, Cat E7300) was used and adapter ligated molecules encoding small RNAs were enriched. Pippin Prep size selection (Sage Science) with 3% agarose was performed according to the following parameters: BP Start = 105 bp, BP End = 155 bp. The cBot was used to apply 25pM of the sequencing library to a HiSeq v4 flowcell (Illumina) and the HiSeq SR Cluster Kit (Illumina, Cat. GD-401-4001) was used for clonal amplification. Following, the flowcell was transferred to the HiSeq 2500 instrument and used in 50 cycles single read sequence run performed with HiSeq SBS Kit v4 reagents (Illumina, Cat. FC-401-4002). Mouse Ensembl gene annotations (build 74) were downloaded and converted to genePred format. Splice junction sequences were generated using USeq’s (v8.8.9) MakeTranscriptome application using a radius of 46. These splice junction sequences were added to the mouse chromosome sequences (mm10) and run through novoindex (v2.8) to create the transcriptome index. Reads were aligned to the transcriptome reference index with Novoalign (v2.08.03), allowing up to 50 alignments for each read. USeq’s SamTranscriptomeParser application was used to select the best alignment for each read and convert the coordinates of reads aligning to splices back to genomic space. Read counts for each mature miRNA (mirBase v21) were generated using USeq’s DefinedRegionDifferentialSeq application. These counts were used in DESeq2 to measure the differential expression between each group.

### Ingenuity pathway analysis

Differentially expressed genes which were identified in Nrf2-TG mice compared with NTG were subjected to Ingenuity Pathway Analysis (IPA^®^) (Qiagen). IPA was ran for Nrf2-TG group relative to the NTG expression profile. Top canonical pathways were identified with the list of genes involved provided the significantly altered biological functions with corresponding disease annotations. Top 30 DEGs were filtered using log2Fold change levels in Nrf2-TG mice within each major biological function category were assembled into heat maps.

### Statistical Analysis

All data are represented as mean ± SEM. Student’s t-test was used for comparisons in NTG vs Nrf2-TG. All analyses were performed using GraphPad Prism 7. Differences were considered significant at the values of *p< 0.05, **p< 0.01, and ***p< 0.001.

## Results and Discussion

### Cardiac Specific Nrf2 Transgenic Mouse Model

PCR for mouse Nrf2 transgene (TG) using primers generated for alpha-MHC promoter (forward) and Nrf2 region (reverse) yielded 350bp for TG expression and product for FABP primers at 250bp in all samples confirming the basal DNA (Fig 1A). Further, semi quantitative PCR using Nrf2 gene specific primer (endogenous) showed increased Nrf2 expression three fold computed using NIH ImageJ compared to NTG mice (Fig 1B). PCR for *Arbp1* primers were used as housekeeping control (Fig 1B). In addition, real time qPCR validation of Nrf2 gene from the 2 experimental groups (n=4-6/group) showed increased expression of endogenous Nrf2 in mNrf2-TG comparison with NTG (Fig 1C). Subsequently, Nrf2-TG mice also exhibited 1.8 fold increases in glutathione levels compared NTG mice (Fig 1D). Hierarchical clustering of genome data of individual samples using next generation RNA sequencing (NGS) data (count normalized FPKM) showed distinctive grouping in accordance with their respective genotypes (Fig 1E). These observations demonstrated a positive role for Nrf2 as a potential transcriptional regulator of myocardial redox state.

**Figure 1.**
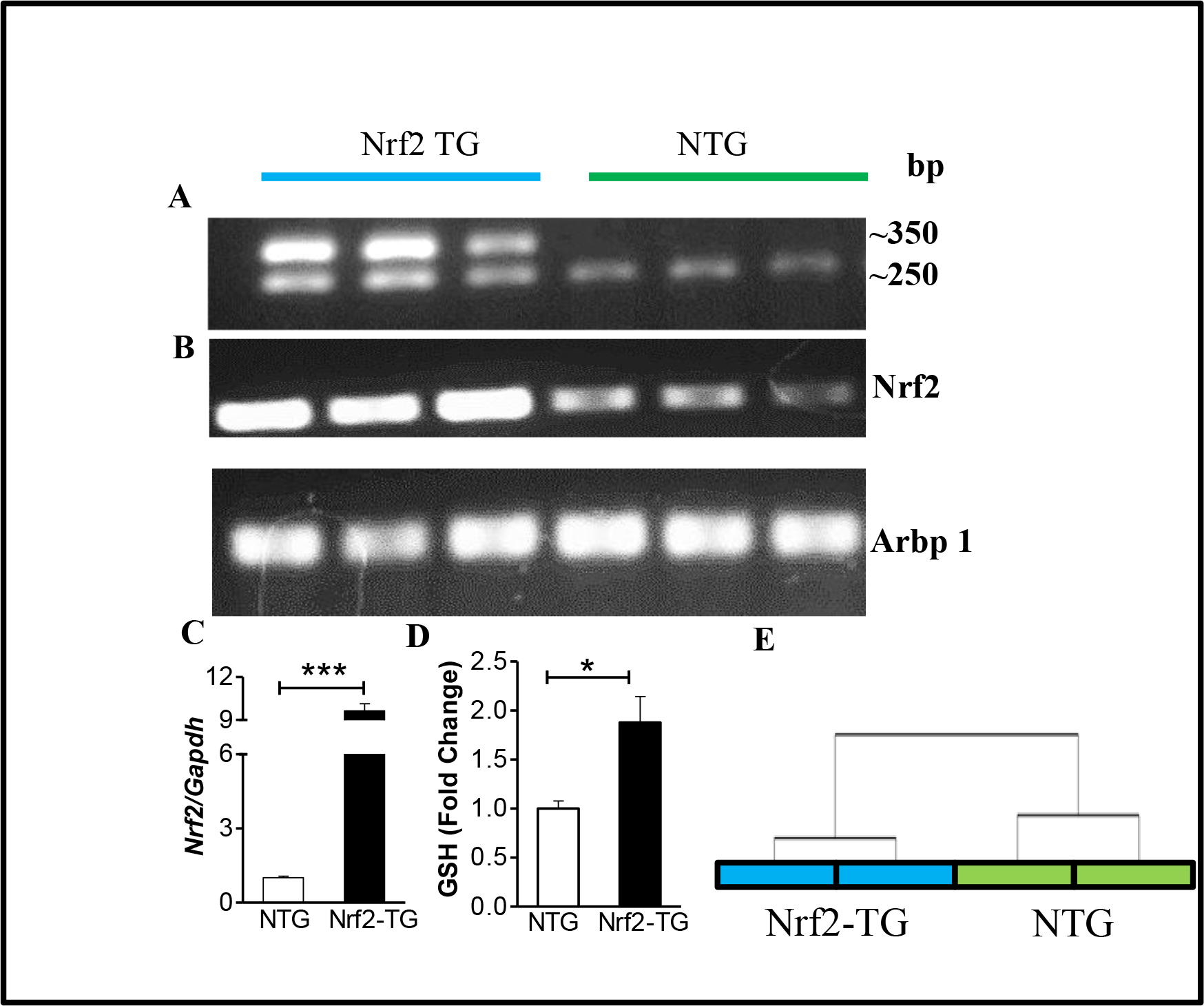
Nrf2 Transgenic mouse model: **A** Genotyping PCR for Nrf2-TG using a-MHC forward and Nrf2-R primers, PCR for FABP primers showed bands at ~250bp used as a DNA control. **B** Semi-quantitative PCR for Nrf2 using gene specific primer showing increased Nrf2 expression in Nrf2-TG compared to NTG. Arbp1 used as control. **C** Real time qPCR for Nrf2 gene using cDNA from hearts of three experimental groups confirms increased Nrf2 expression in Nrf2 TG compared to NTG mice(n=4-6/group), ***p<0.001. **D** Nrf2-TG mice showed increased GSH levels compared to NTG, *p<0.05. **E** Hierarchical clustering of normalized FPKM values from RNAseq revealing distinctive grouping of at least 2 samples in each group.

### Cardiac Transcriptome signature in the mNrf2-Transgenic mouse

The high throughput sequencing data of myocardial RNA from mNrf2-TG mice detected 12,301 mRNAs. Primary analysis identified that 46% (5727 genes) of the total transcriptome showed a noticeable changes in gene expression (log2 FC >1.5) compared to NTG, establishing them as differentially expressed genes (DEGs). Within those DEGs, about 21.6% were significantly altered with log2 FC more than 2 (Fig 2A). Majority of these DEG’s were found to be upregulated in mNrf2-TG hearts as illustrated in the heat map (Fig 2B) displaying genes that are noticeably altered with log2 FC of ≥1.5. However, in the cases where exceptional changes were identified in 89 genes (or 0.7 % of detected transcriptome) with log2 FC >5, the number of downregulated DEGs were twice (n=57) compared to upregulated ones (n=31). Top expression changes were detected in *Anks6, Dennd4a, Cxcl14, Gsta2, Gsta3, Gclm, Gstm1 and Map1a* which were the most upregulated in response to the transgene.

**Figure 2.**
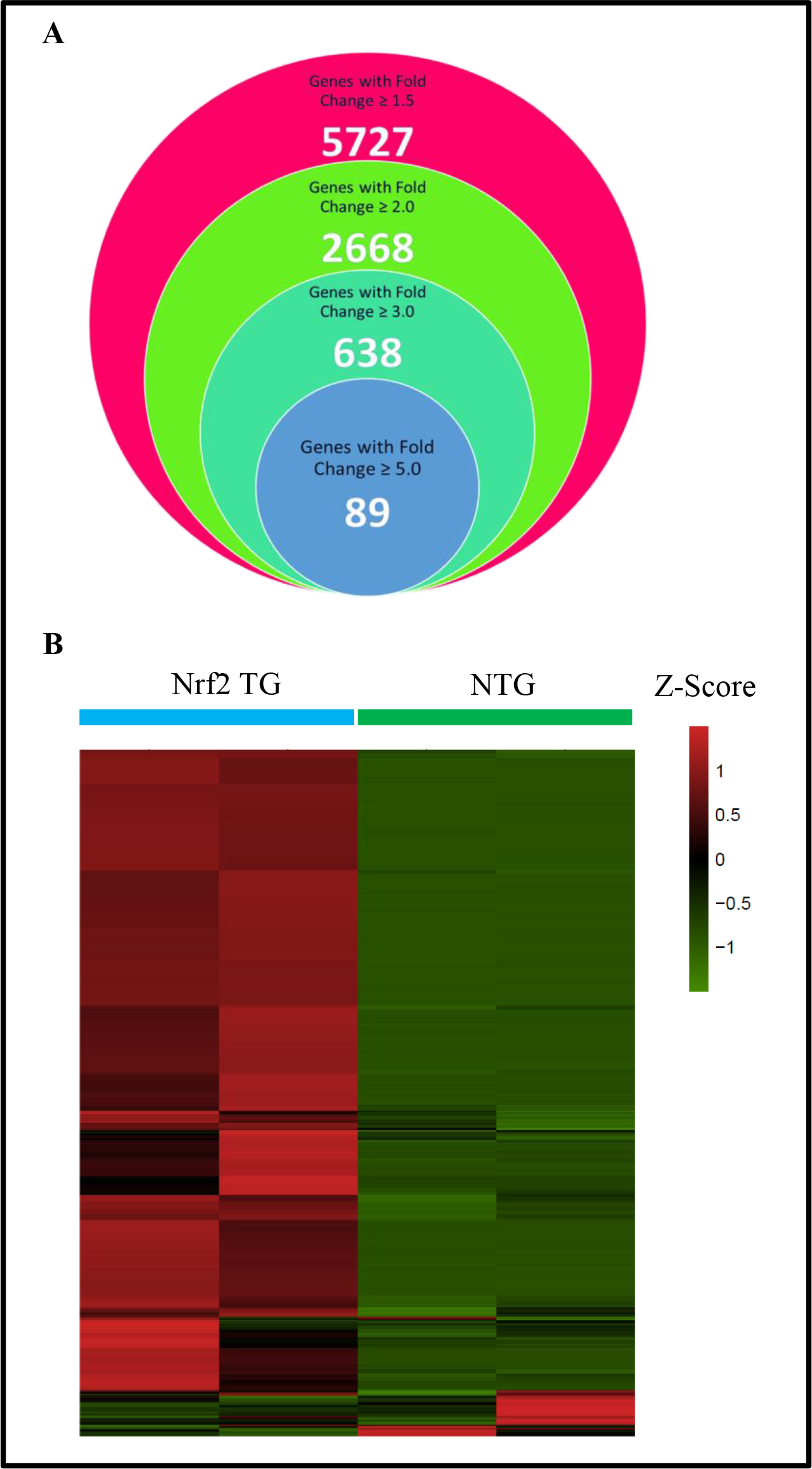
Transcriptomic changes in Nrf2-transgenic vs Non-transgenic mice: **A.** Venn diagram demonstrating the number of differentially expressed genes from NGS RNAseq in Nrf2 TG compared to NTG mice and subsets of significantly altered transcriptomes with fold changes ranging from 1.5 to more than 5. **B** Heat map representation of 5727 differentially expressed genes (of more than 1.5 FC) in Nrf2 G hearts compared to NTG.

### Top Canonical pathways identified and their biological functions in mNrf2-Transgenic mouse

Differentially expressed genes were processed by functional analysis using Ingenuity Pathway Analysis (IPA) and the top pathways enriched in mNrf2-TG versus NTG on biological processes were expectedly related to Nrf2-mediated redox mechanisms and associated pathways (Fig 3A). The topmost pathway that was altered in Nrf2 Transgenic mice was Nrf2 mediated oxidative stress response pathway. The other significantly altered pathways identified were ILK Signaling, hypoxia signaling in the cardiovascular system, glutathione-mediated detoxification, and cardiac hypertrophy signaling (Fig. 3). From a total of 4 out of 10 most altered pathways collectively insisted that the profile of Nrf2-TG hearts is displaying a distinct redox transcriptome profile than NTG mice (Fig 3B).

**Figure 3.**
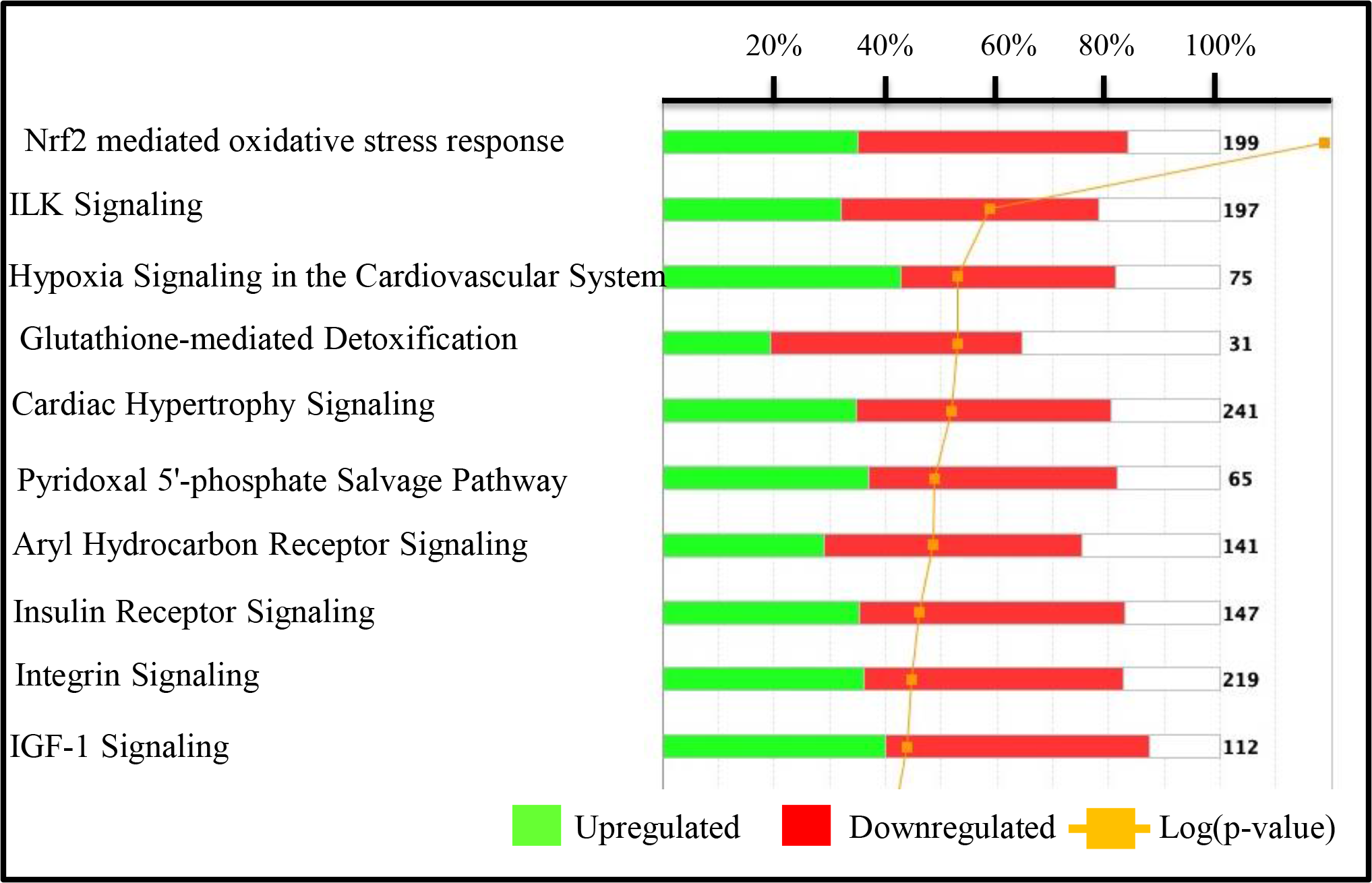
Top 10 Canonical pathways that are potentially regulated by the genes identified in Nrf2 Transgenic hearts. Numbers to the right of the bars indicate the total number of genes contributing the pathway

### Effect of Nrf2 transgene on Canonical pathway associated with Nrf2-glutathione mediated stress response and cardiac hypoxia signaling in mouse myocardium

Identification and profiling of top most altered canonical pathways were made possible by Ingenuity Pathway Analysis (IPA) using mRNAseq data from Nrf2-TG and NTG hearts. As anticipated, the top most altered canonical pathways were Nrf2 mediated stress response and Glutathione mediated detoxifications, which were prominently augmented into a combined representation as they were construed to be objectively homologous from a redox perspective. Out of 230 genes known to constitute these pathways, about 186 were found to be present in the detected transcriptome and 48 DEGs amongst them had a significant log2 fold change >2. The heat map (Fig. 4A) illustrates the top 30 DEGs in the aforementioned pathway. *Acta2*, as encoded protein is known to be involved in vascular contractility and blood pressure homeostasis was found be the most upregulated (Log2 fold change of 6.4) in this pathway. Accompanying, with similar levels of fold change was noticed among *Nfe2l2, Kl (Klotho), Txnrd1 and Gst* family of detoxification enzymes. qPCR validation(Fig 3C) for *Acta2* (23) and *Txnrd1* among others confirmed direction of expression changes revealed by NGS sequencing. Other Validated mRNAs were *Gsta3, Nqo1* which were remarkably upregulated in Nrf2-TG mice and GSR which was appreciably upregulated in the same. Cardiac hypoxia signaling pathway was also found to be one among the top 4 altered pathways. In this particular pathway 60 mRNAs were detected in Nrf2-TG mice out of 75 genes that constitute this pathway and few of the mRNAs (16) were found to significant levels of fold change(Log2 FC >2). These genes include *Nqo1* (log2 FC of 3.4, up), *Vegfa*, *Cdc34*, *Jun*, *Ikbkg*, ubiquitin conjugating enzymes (*Ube2O*, *Ube2I*) and most importantly *Hif1A*, the master transcriptional regulator of cellular and developmental response to hypoxia. qPCR was performed to confirm the consistency of NGS data and the expression of *Crebbp, Atf2* and *Ube2M* were confirmed to be upregulated and *Hif1A* was confirmed to be downregulated (Fig. 4B).

**Figure 4.**
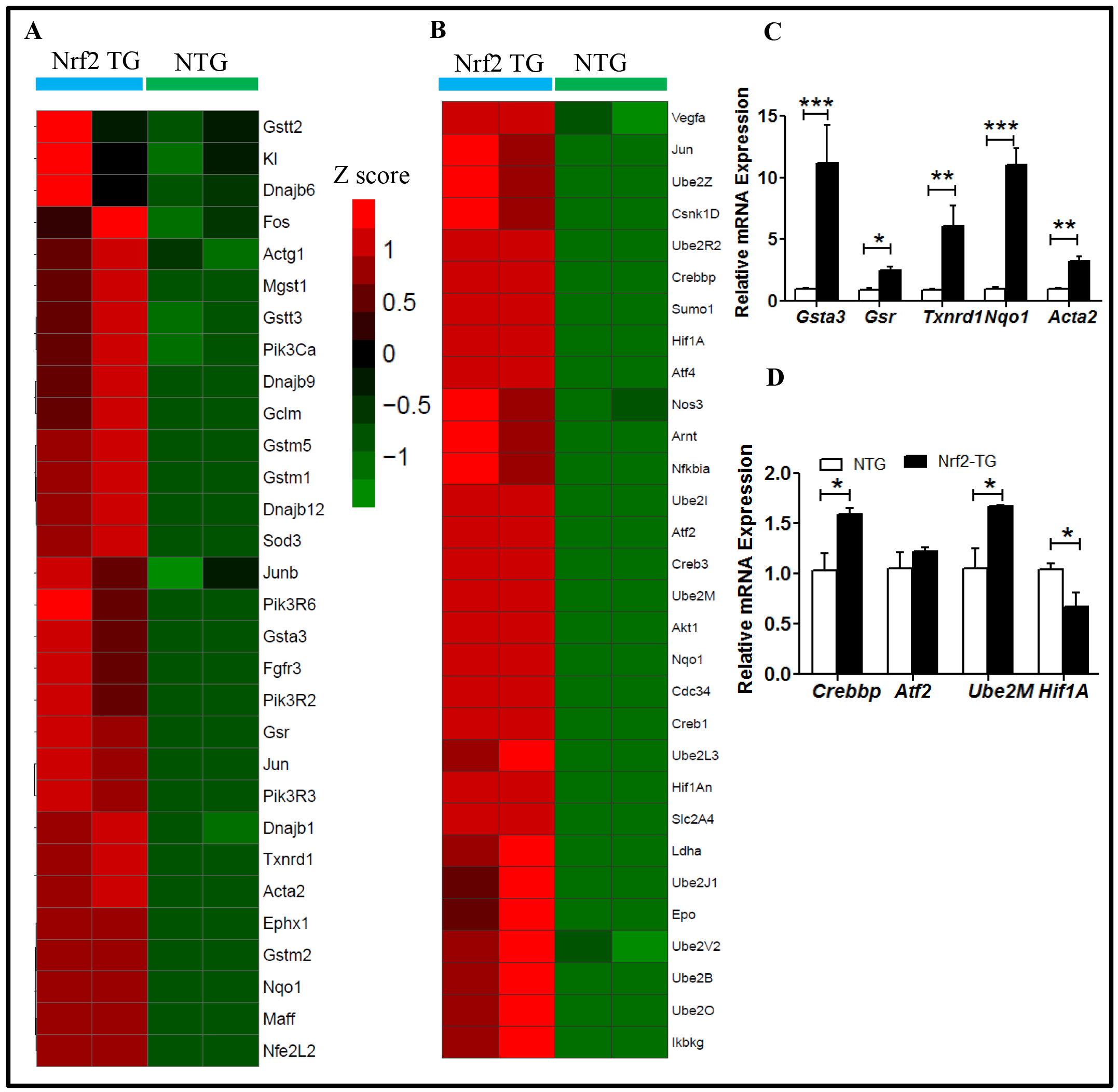
Significantly altered Canonical pathways with biological relevance: **A and B** Heat map illustrating NGS-RNAseq normalized FPKM values of top 30 DEGs in Nrf2-Glutathione mediated oxidative stress response/detoxification pathway and Hypoxia signaling in cardiovascular system pathway respectively identified by Ingenuity^®^ Pathway Analysis (IPA^®^). **C** Red hue of Z-score indicates upregulation and green hue indicates downregulation. Dark tinge of either colors and black indicates insignificant or no change in gene expression. **C,D** Real time qPCR validation of 4-5 genes from their respective pathways of Nrf2-Glutathione mediated oxidative stress response and Hypoxia signaling pathways validated the altered expression compared between Nrf2 TG vs NTG groups. (n=4-6/group).*p<0.05, **p<0.001, ***p<0.0001.

### Effect of Nrf2 transgene on Integrin-Linked Kinase (ILK) and Cardiac hypertrophy signaling

Molecular components of the cardiac mechanical stretch sensing are also appear to undergo significant changes from ILK signaling in the heart. In this specific pathway, about 37/152 detected mRNAs transcripts have undergone significant expression changes (Fig 5A). It is quite uncertain that these changes are in respect to remodeling to retain the mechanical homeostasis of heart function. The heat map illustrates the top 30 genes represented in this pathway (Fig 5A). *KL* and *Acta2* which are present in the oxidative stress response pathway also exists in the ILK pathway. *Acta2*’s differential was validated through qPCR (Fig 4A). Myl9 that encodes for myosin light chain that regulates muscle contraction has also been found to be the most altered in this pathway. This was established through quantitative PCR (Fig 5A) which also showed significant upregulation of *Myl9* in Nrf2-TG mice relative to NTG. In addition randomly picked genes for validation included *Mtor, Itgb5* and *Pik3ca* were substantiated the sequencing results (Fig 5C). Another responsive action to counteract this would be any changes seen in cardiac hypertrophy signaling (Fig 5B). About 1/3 of the mRNAs have undergone significant expression changes and *Myl9* that encodes for myosin light chain that regulates muscle contraction has been found to be the most altered in this pathway. Genes were selected at random to reaffirm the NGS data through qPCR and it was established that *Map3k7, Map3k10, srf1* and *Tgfbr2* were upregulated and *Map3k6* was downregulated in Nrf2-TG mice (Fig 5D).

### Micro-RNA expression profile in the Nrf2 transgenic hearts

Since micro RNAs serve as regulators of certain host of mRNA expression (24) at post transcriptional level, the Nrf2 transgene-dependent changes in miRNA expression were studied in parallel using the data obtained from miRNAs sequencing (25). In a total of 556 transcripts, about 26 (or about 4.6%) miRNAs had significantly altered levels of expression in response to Nrf2 transgene. Among the 26 miRNAs, 15 were upregulated and 11 were downregulated significantly in the myocardium of Nrf2-TG mice (26). The represented heat map (Fig 6B) illustrates the directions of expression change in the miRNA transcriptome of Nrf2-TG mice. Heat map of top 26 miRNAs expression changes with significant differential expression (Log2 FC >2) are illustrated (Fig 6C). Reliable changes (Log2 FC >1.5) were noted in miR-451a, miR-144-3p, miR-3068-3p, miR-671-5p, miR-582-3p. These were confirmed by qPCR validation for miR-671-5p, miR-671-3p, 677-5p, miR-361-5p, miR-5099, miR-3535, miR-155-3P, miR-144-3p

**Figure 5.**
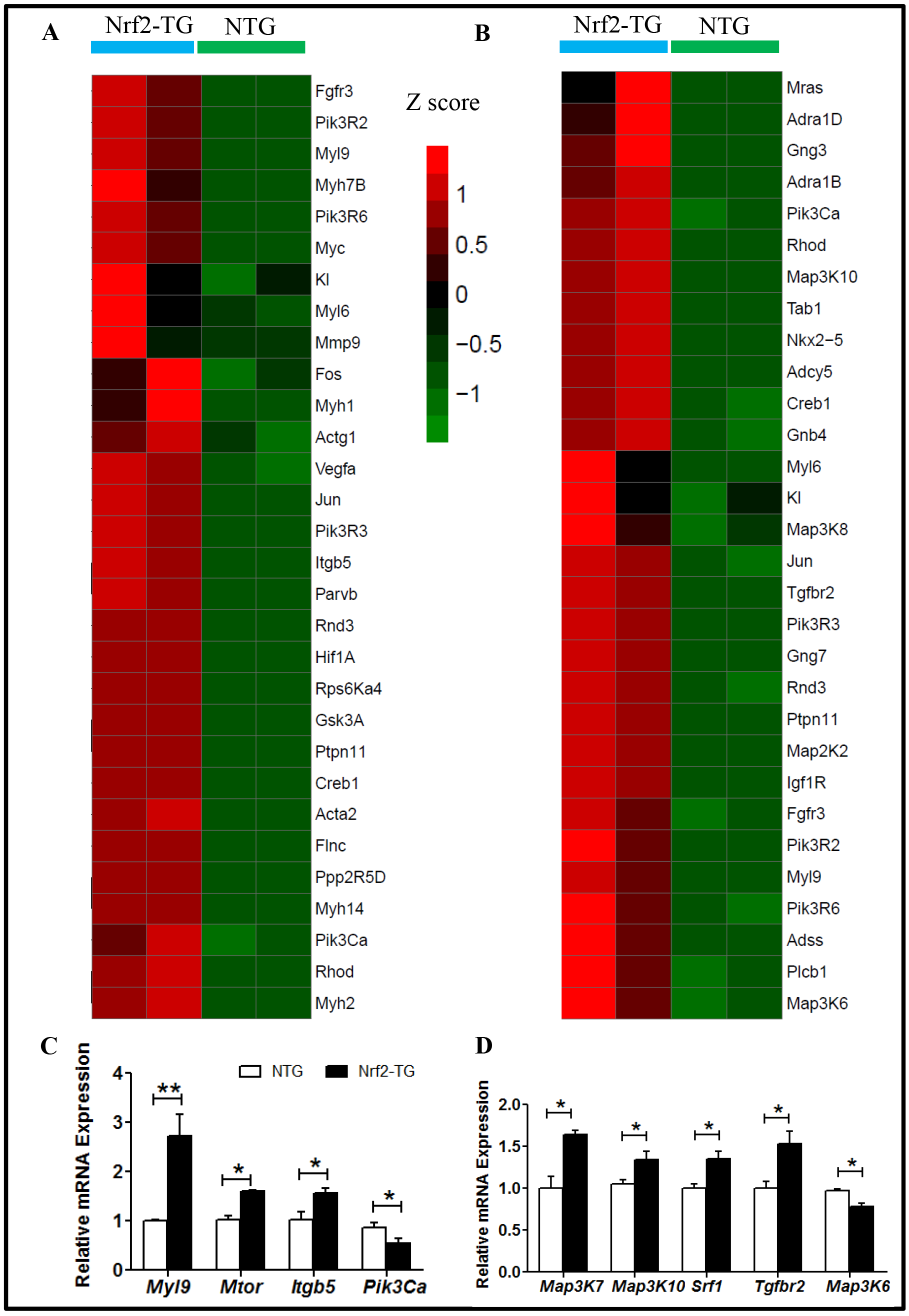
Most altered Canonical pathways with biological relevance: **A,B** Heat map illustrating NGS-RNAseq normalized FPKM values of top 30 DEGs in ILK signaling and Cardiac hypertrophy signaling respectively identified by IPA^®^. **C,D** qPCR validation of 4-5 genes with their respective pathways of ILK signaling and Cardiac hypertrophy signaling validated the altered expression between Nrf2 TG and NTG groups. (n=4-6/group). *p<0.05, **p<0.001, ***p<0.0001.

**Figure 6.**
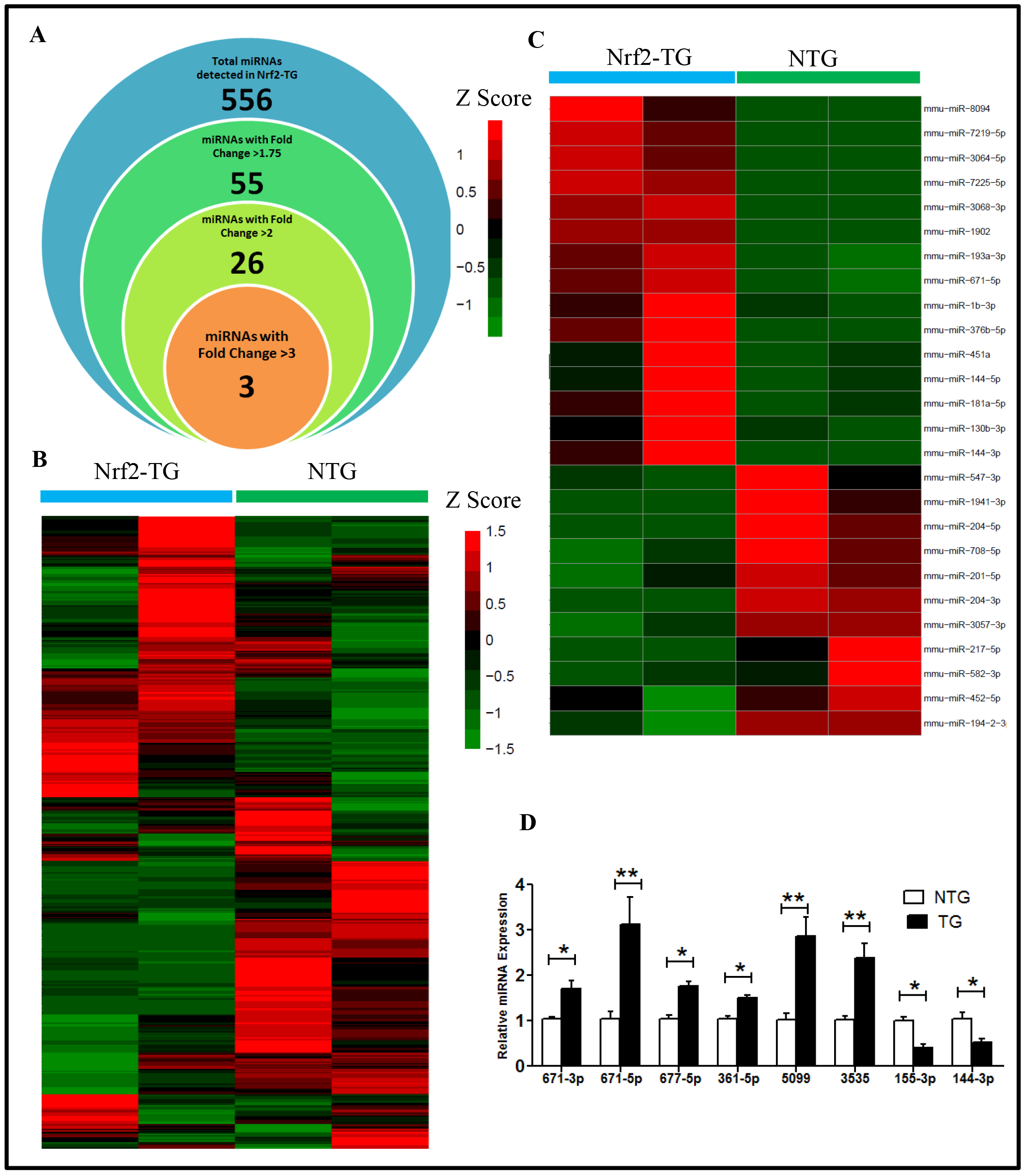
Nrf2 mediated regulation of miRNAs expression in Nrf2-TG myocardium: **A** Venn diagram representing the significantly altered number of miRNAs detected in myocardium from small-RNA seq data with subsets of miRNAs defined by their fold change ranging from 1.75 to more than 5. **B** Heat map for all detected miRNAs (n=556) using normalized FPKM from NGS-small RNA seq in the myocardium of Nrf2 TG vs NTG illustrating sets of upregulated and downregulated miRNAs between them. **C** Heat map from miRNA seq derived FPKM illustrating most differentially expressed miRNAs with fold change of 2 or more between Nrf2 TG and NTG. **D** qPCR validation of 8 miRNAs which have high differential expression between Nrf2-TG and NTG groups. (n=4-6/group). *p<0.05, **p<0.001, ***p<0.0001.

## Conclusion

NGS results and IPA revealed that excess Nrf2 in the myocardium significantly altered the transcriptional (RNA) and post-transcriptional (miRNA) targets, even under unstressed conditions. Further, our unbiased approach/methods employed in the study has depicted diverse changes in both mRNA and miRNA expression, which impacted on various pathways that involve in key biological process/signaling. The top most pathways or biological functions that are regulated by excess Nrf2 is redox regulation, which includes 1) Nrf2 mediated oxidative stress, 2) glutathione mediated detoxification, 3) hypoxia signaling and 4) cardiac hypertrophy signaling. However, experimental validation using luciferase based approach is required for the newly reported genes to specify that these genes are regulated by Nrf2. Further, future studies will be focused on developing specific miRNA/mRNA knockout and transgenic *in vitro* and/or *in vivo* models to study the functional mechanism of a particular gene or miRNA. One limitation of our study is that only male mice were used for sequencing; however qPCR validation has been performed in both genders.

## Acknowledgements

This study was supported by funding from NHLBI (HL118067), NIA (AG042860), the AHA (BGIA 0865015F), the Division of Cardiovascular Medicine/Department of Medicine, University of Utah and the start-up funds by Departments of Pathology and Medicine, the University of Alabama at Birmingham, AL.

The authors would like to thank Dr. Brian Dalley, High-Throughput Genomics and Bioinformatic Analysis Shared Resource of the Huntsman Cancer Institute at the University of Utah for their assistance with RNA sequencing experiments and data analyses. Authors’ also thank Ms. Jennifer Hong and Ms. Nancy Atieno for their assistance in maintaining the animal colonies and Mrs. Jennifer Schroff for editorial assistance.

